# Biocalcification in porcelaneous foraminifera

**DOI:** 10.1101/2023.10.02.560476

**Authors:** Zofia Dubicka, Jarosław Tyszka, Agnieszka Pałczyńska, Michelle Höhne, Jelle Bijma, Max Janse, Nienke Klerks, Ulf Bickmeyer

**Affiliations:** Ecological Chemistry, Alfred-Wegener-Institut Helmholtz-Zentrum für Polar-und Meeresforschung, Bremerhaven D-27570, Germany; GFZ German Research Centre for Geosciences, Telegrafenberg, 14473 Potsdam, Germany; Faculty of Geology, University of Warsaw, Warsaw PL 02-089, Poland; Research Centre in Kraków, Institute of Geological Sciences, Polish Academy of Sciences, Kraków 31-002, Poland; Marine Biogeosciences, Alfred-Wegener-Institut Helmholtz-Zentrum für Polar-und Meeresforschung, Bremerhaven D-27570, Germany; Burgers’ Ocean, Royal Burgers’ Zoo, Arnhem 6816 SH, The Netherlands

**Keywords:** amorphous calcium carbonate, mesocrystals, biomineralization, fluorescence, Paleozoic biocalcification, evolution

## Abstract

Living organisms control the formation of mineral skeletons and other structures through biomineralization. Major phylogenetic groups usually consistently follow a single biomineralization pathway. Foraminifera, which are very efficient marine calcifiers, making a substantial contribution to global carbonate production and global carbon sequestration, are regarded as an exception. This phylum has been commonly thought to follow two contrasting models of either *in situ* “mineralization of extracellular matrix” attributed to hyaline rotaliid shells, or “mineralization within intracellular vesicles” attributed to porcelaneous miliolid shells. Our previous results on rotaliids along with those on miliolids in this paper question such a wide divergence of biomineralization pathways within the same phylum of Foraminifera. We found that both groups produced calcareous shells via the intravesicular formation of unstable mineral precursors (Mg-rich amorphous calcium carbonates) supplied by endocytosed seawater and deposited at the site of new wall formation within the organic matrix. Precipitation of high-Mg calcitic mesocrystals took place *in situ* and formed a dense, chaotic meshwork of needle-like crystallites. We did not observe deposition of calcified needles that had already precipitated in the transported vesicles, which challenges the previous model of miliolid mineralization. Hence, Foraminifera utilize less divergent calcification pathways, following the recently discovered biomineralization principles. Mesocrystalline chamber walls are therefore apparently created by accumulating and assembling particles of pre-formed liquid amorphous mineral phase within the extracellular organic matrix enclosed in a biologically controlled privileged space by active pseudopodial structures. Both calcification pathways evolved independently in the Paleozoic and are well-conserved in two clades that represent different chamber formation modes.

## Introduction

Over the past 500 million years, living organisms evolved different skeleton crystallization pathways. Very popular in nature is the mineralization of the extracellular matrix, for example, in crustacean cuticles, mollusk shells, vertebrate bones, and teeth composed of dentin and enamel (Weiner and Addadi, 2011; Kahil et al., 2021). Radial foraminifera represented by rotaliids have been traditionally interpreted to make use of this crystallization mode (Weiner and Addadi, 2011).The other two pathways are intravesicular and are characterized by either production of amorphous unstable phase within a large vesicle, such as a syncytium, well documented for sea urchin larvae (Beniash et al., 1997) or crystallization of calcite elements within smaller vesicles located in the intracellular space, as seen in fish that form guanine crystals and coccolithophores to produce coccoliths (Weiner and Addadi, 2011; Kahil et al., 2021). This model has also been attributed to the formation of porcelaneous shells by miliolid foraminifera (Weiner and Addadi, 2011) based on the model proposed by Berthold (1976) and followed by Hemleben et al. (1986). As such, mineralization of shells in Foraminifera is believed to follow two highly contrasting pathways. The current theory states that Miliolida, characterized by imperforate, opaque milky-white shell walls (porcelaneous) (Angell, 1980; Hemleben et al. 1986; de Nooijer et al., 2009), produce fibrillar crystallites composed of Mg-rich calcite within tiny vesicles enclosed by cytoplasm. Miliolid shells are made of randomly distributed calcite needles that form a dense meshwork of chaotic crystallites that cause light reflection, resulting in opaque (porcelaneous) milky walls (Hohenegger, 2009). Calcite needles are thought to be precipitated completely within these vesicles and then transported to the site of chamber formation to be released via exocytosis (Berthold, 1976; Hemleben et al. 1986; Angell, 1980; de Nooijer et al., 2008, 2009).The pre-formed needles or needle stacks are believed to be continuously embedded in an organic matrix in the shape of the new chamber until the wall is completed. Although this model is commonly accepted, it has never been sufficiently documented *in vivo*, and it does not resolve several conflicting issues. First of all, the question is how pre-formed bundles of parallel calcitic needles are transformed into randomly oriented needles within the shell wall. It is difficult to explain, if there is no recrystallization process within the wall structure after discharging the calcite crystallites. This problem was already emphasized by Hemleben et al. (1986). Secondly, why the newly constructed wall is still translucent after deposition of random crystals. We would expect a thin milky opaque layer of the new wall under normal transmitted light, as well as polarized crystals of calcite under crossed nicols. Angell (1980) on his plate 2 presenting porcelaneous chamber formation in miliolid *Spiroloculina hyalina* Schulze clearly documented the polarization front being shifted circa a half of the length of the new chamber behind the leading edge of the forming chamber. This shift represented more than an hour. Therefore, polarization was missing in the early and middle stage of chamber formation. It means that Angell’s (1980) time lapse micrographs of the chamber formation were in conflict with the imaging under TEM. It seems that Angell (1980) was aware of that problem and stressed that calcification had to be “intense enough to show under crossed nicols lags behind the leading edge of the forming chamber” (p. 93, pl. 2 fig. 12/caption). In fact, all experiments that show the “crystal vacuoles” (sensu Angell, 1980) documented under TEM (Berthold, 1976; Hemleben et al. 1986; Angell, 1980) required fixation of the samples, which was prone to post-fixation artifacts of unwanted calcite precipitation.

Our goal is to test whether the miliolid shell is produced by “agglutination” of premade needle-like calcitic crystallites, and in consequence, whether this large group of calcareous Foraminifera follow crystallization within smaller vesicles located in the intracellular space. Therefore, we re-examined the mineralization process in Miliolida based on experiments on a living species, *Pseudolachlanella eburnea* (d’Orbigny) (Fig. 1). This taxon was selected to facilitate replicated observations of chamber growth under controlled culture conditions. We included observations of *in vivo* biomineralization using multiphoton and confocal laser scanning microscopy (CLSM) followed by analyses of fixed specimens at different stages of chamber formation by high-resolution field emission scanning electron microscopy (FE-SEM) coupled with energy dispersive X-ray spectrometry (EDS). Our new FE-SEM data challenge the current understanding of the biomineralization of miliolid foraminifera and such a significant divergence of biomineralization pathways within the Foraminifera.

**Figure 1.**
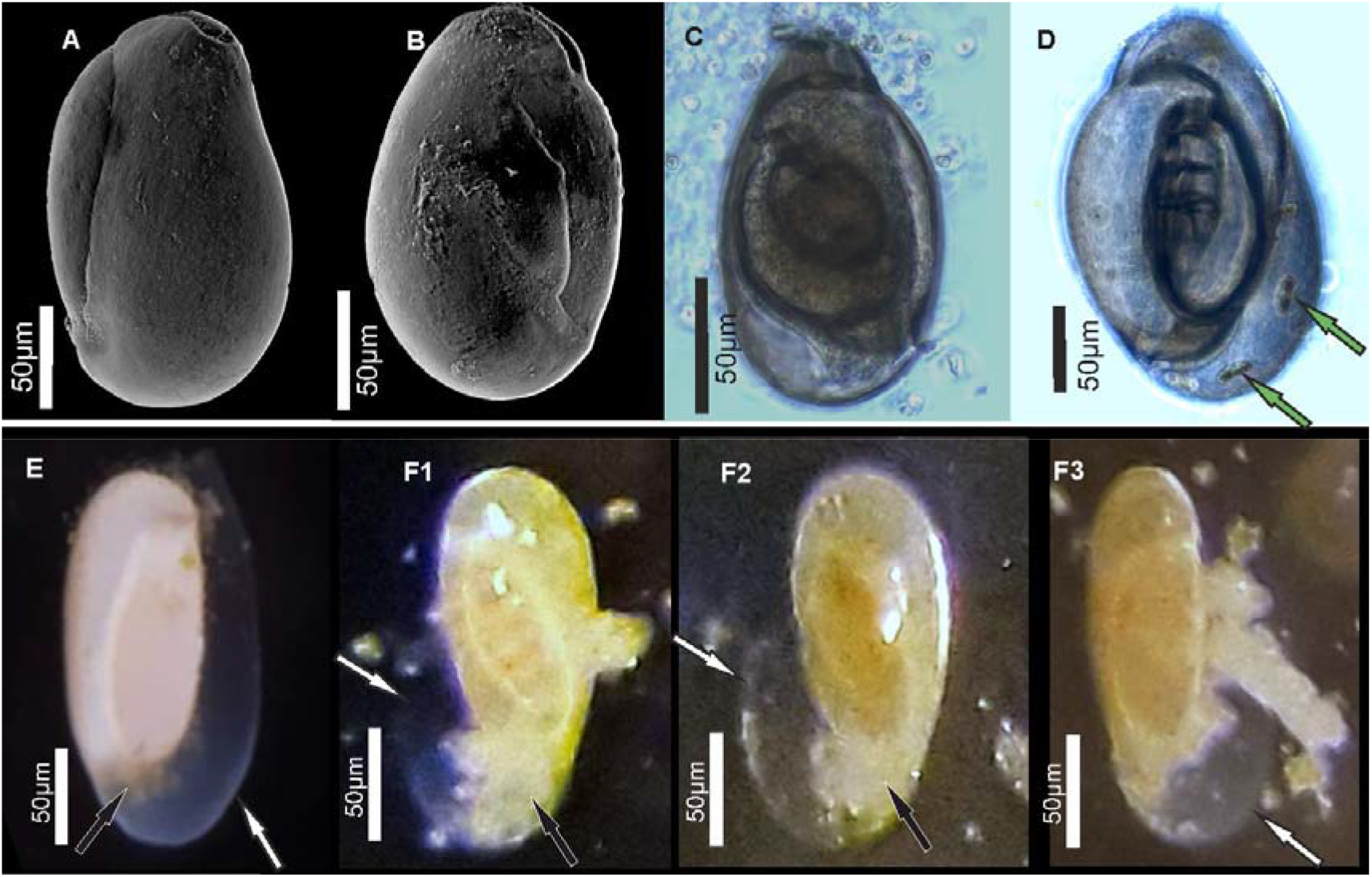
Specimens of miliolid foraminifera, identified as *Pseudolachlanella eburnea* (d’Orbigny), used for experimental studies. (A, B) SEM, (C, D) transmitted light microscope, and (E, F) stereomicroscope images. White arrows show the outer organic sheath of a new chamber during its gradual calcification expressed by its gradual appearance from complete transparency to milky and opaqueaspect (E, F). Black arrows indicate a small mass of cytoplasm extruded from the aperture of exiting the chamber. Green arrows point to incorporated algae.

## Results

All replicated *in vivo* experiments on *Pseudolachlanella eburnea* facilitated by CLSM imaging with the application of membrane impermeable Calcein and FM1-43 membrane dyes (performed in separate experiments) showed intravesicular fluorescence signals from groups of moving vesicles (1–5 μm in size) inside the cytosol (Fig. 2A, B, Movies S1 and S2). The fluorescent vesicles inside the cytosol contained seawater, as documented by fluorescence of membrane impermeable Calcein. These vesicles were taken up by endocytosis indicated by FM1-43 staining. This dye stains the cell membranes and indicates all endocytic vesicles by fluorescence, whereas the other intracellular vesicles remain unstained (Amaral et al., 2011). Both dyes demonstrate the uptake of seawater via the endocytosis of vesicles that are approximately 1–4 μm in diameter and move through the entire cell.

**Figure 2.**
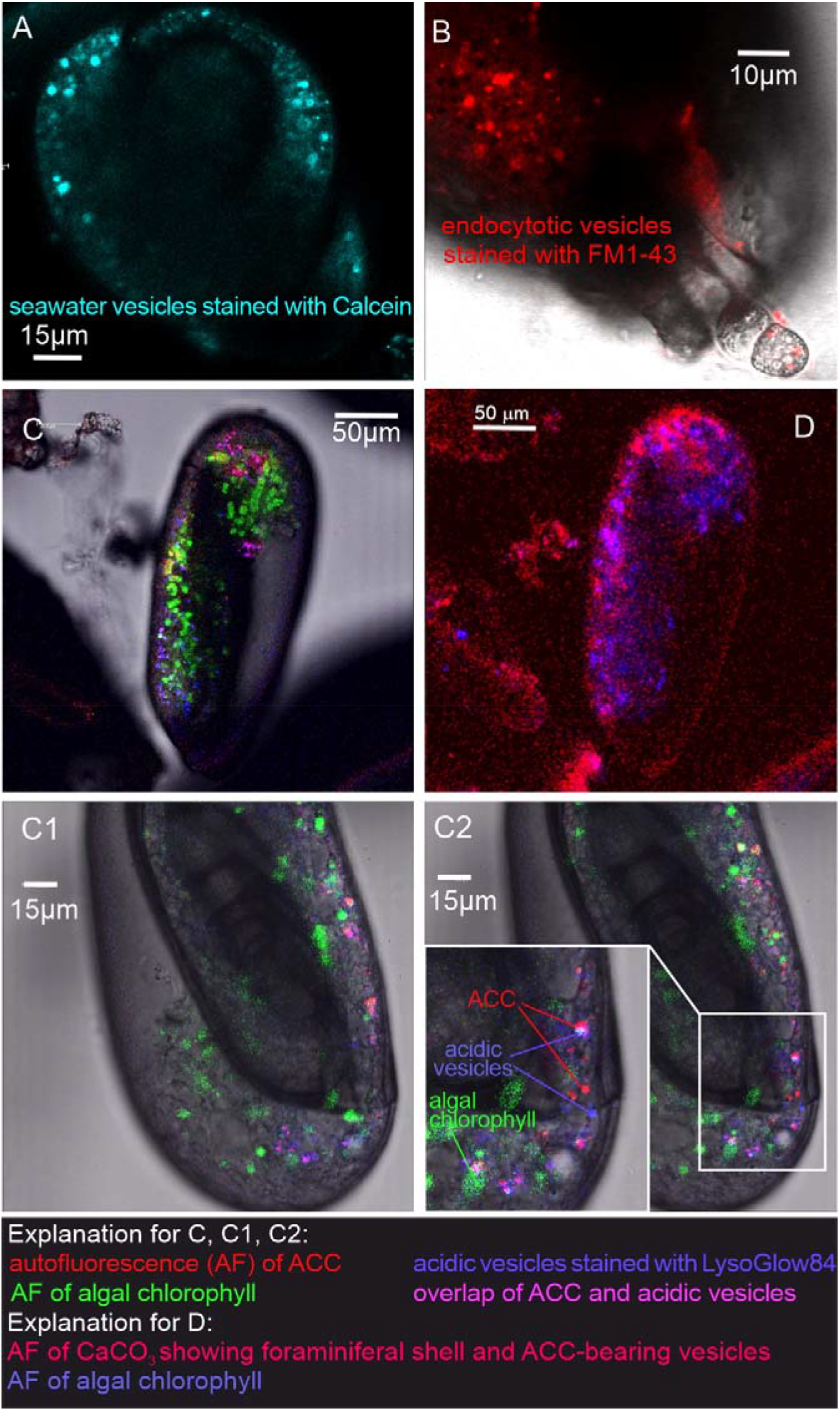
Fluorescence images of living *P. eburnea* conducted by Laser Scanning Microscopy. A - Cell impermeable Calcein (cyan) indicating endocytotic seawater vesicles, see Movie 1. B - FM1-43 membrane dye indicating endocytotic vesicles (red), see Movie 2. C, E - LysoGlow84 indicating acidic vesicles (navy blue), autofluorescence of chloroplasts (green), and Mg-ACC pools (red), see Movies 3 and 4, (note the overlap of ACC and acidic vesicles is marked in lilac). D - Autofluorescence image with reduced threshold of the studied Miliolida species (exc. 405 nm) showing algal chlorophyll (blue) and CaCO_3_ (red), both ACC and calcite shell.

Additional LysoGlow84 staining revealed numerous acidic vesicles in the cytosol the presence of (Fig. 2C, Movies S3 and S4). Acidic vesicles were accompanied by other vesicles (approximately 1–2 μm in size) that show autofluorescence upon multiphoton excitation at 405 nm (emission 420–480 nm), shown in red in Figure 2C. This wavelength partly permeates the shell to excite autofluorescence interpreted as associated with ACCs (see Dubicka et al., 2022). The autofluorescence of the shell itself is also present (Figure 2D), however, it is not clearly visible because the fluorescence of ACCs is much stronger. The intensity of the laser light is reduced because the multiphoton light has to pass through a thick three-dimensional carbonate wall of the foraminiferal shell. Further experimental studies are needed to confirm the ACC source of this autofluorescence and thus definitively eliminate potential organic sources of AF emissions.

In addition, typical chlorophyll autofluorescence (excitation at 405 or 633 nm, emission 650-700 nm, Fig. 2C, Movies S3 and S4 highlighted in green) was detected, which indicated the presence of chloroplasts in microalgae cells. These algal cells have been found to move within the cytosol of the observed specimens, in proximity of acidic vesicles and vesicles characterized by autofluorescence upon UV light (ex. 405 nm). These algal cells may represent facultative endosymbionts, as they were observed only during the chamber mineralization process in specimens with carbonate-bearing vesicles detected by *in vivo* CLSM experiments. They were documented just below the organic matrix (OM) of the newly formed chamber, as seen in the FE-SEM observations as well as just below the organic matrix (OM) of the newly created chamber as seen in the FE-SEM observations (Figs. S1G, and S1H).

Specimens of *P. eburnea*, which displayed vesicles showing autofluorescence under UV light inside the cytosol, were fixed using Method B (see Materials and Methods) coated with a few nanometers of carbon and analyzed by SEM-EDS. The main elements detected in the area of the fixed cytoplasm (Fig. S4) were C, O, Na, Mg, P, S, Cl, K, and Ca (of particular interest were the high contents of Mg and Ca), whereas the main elements detected within the area of the new chamber in the form of a gel-like matter filled with dispersed nanograins were C, O, Na, Mg, S, Cl, and Ca (Fig. S4). The shell content was strongly enriched with Ca relative to the cytoplasm, which showed a much higher Mg/Ca ratio.

FE-SEM observations of the fully mineralized test walls displayed the porcelaneous structures (see Parker, 2017; Dubicka et al., 2018), which are made of three mineralized zones, i.e. (I) extrados that represents an outer mineralized surface (approximately 200–300 nm in thickness; Figs. S1C and S2C); (II) porcelain that denotes the main body of the wall constructed from randomly oriented needle-shaped crystals (up to 1–2 μm in length and approximately 0.2 μm in width). No gel-like matter was observed between the needles of the porcelain structures that appeared in the early stages of wall formation (Figs. 3E; 3E1; S2C, and S3A); and (III) intrados that represents an inner mineralized surface (approximately 200–300 nm in thickness) made of needle-shaped crystallites (Figs. 3E, 3E1 and S1A).

**Figure 3.**
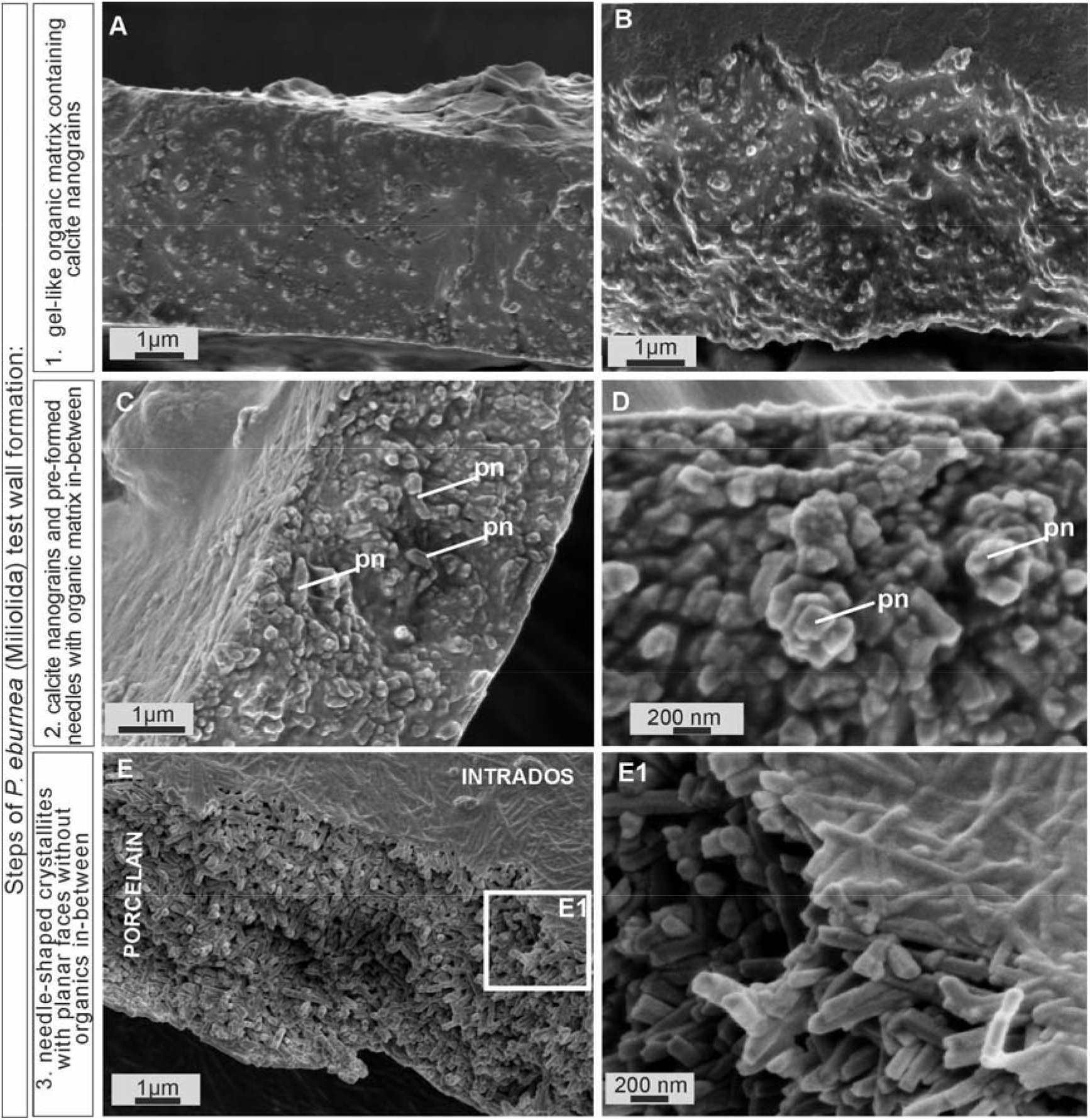
SEM images of the major steps of the formation of *P. eburnea* shell-building components. Test cross-section showing: (A, B) carbonate nanograins within organic matrix, (C, D) nanograins merging into needle-like mesocrystals, (E) fully developed needle-shaped elements; pn –nanograins partly transformed to short needles.

Growing chambers, captured at the various successive stages of chamber formation in different specimens, have revealed the following morphological features: (A) a solitary, thin organic sheath (approximately 200–300 nm thick) that represents the most distal part of the new chamber and is anchored to the older, underlying solid calcified chamber (Fig. 4A); (B) a solitary, outer organic sheath (OOS) filled with spread calcifying nanograins (Figs. 4B; S2A, and S2B); (C) a gel-like matter (4–5 μm in thickness) with a granular texture, bounded on two sides by intrados and extrados, and containing relatively widely spaced, randomly dispersed carbonate nanograins (Figs. 3A-B; 4C; S1A-D); (D) the test inside made of chaotic meshwork of carbonate nanograins partly transformed to short needles with a small amount of gel-like OM in-between (Figs. 3C, D, and 4D); (E) the test inside composed of needle-shaped crystals with planar faces and no apparent remaining gel-like matter (Figs. 3E, E1 and 4E). Carbonate nanograins at the shell construction site were well documented in our SEM-EDS studies (Fig. S4). Both fixation methods (see Material and Methods) yielded highly consistent results.

**Figure 4.**
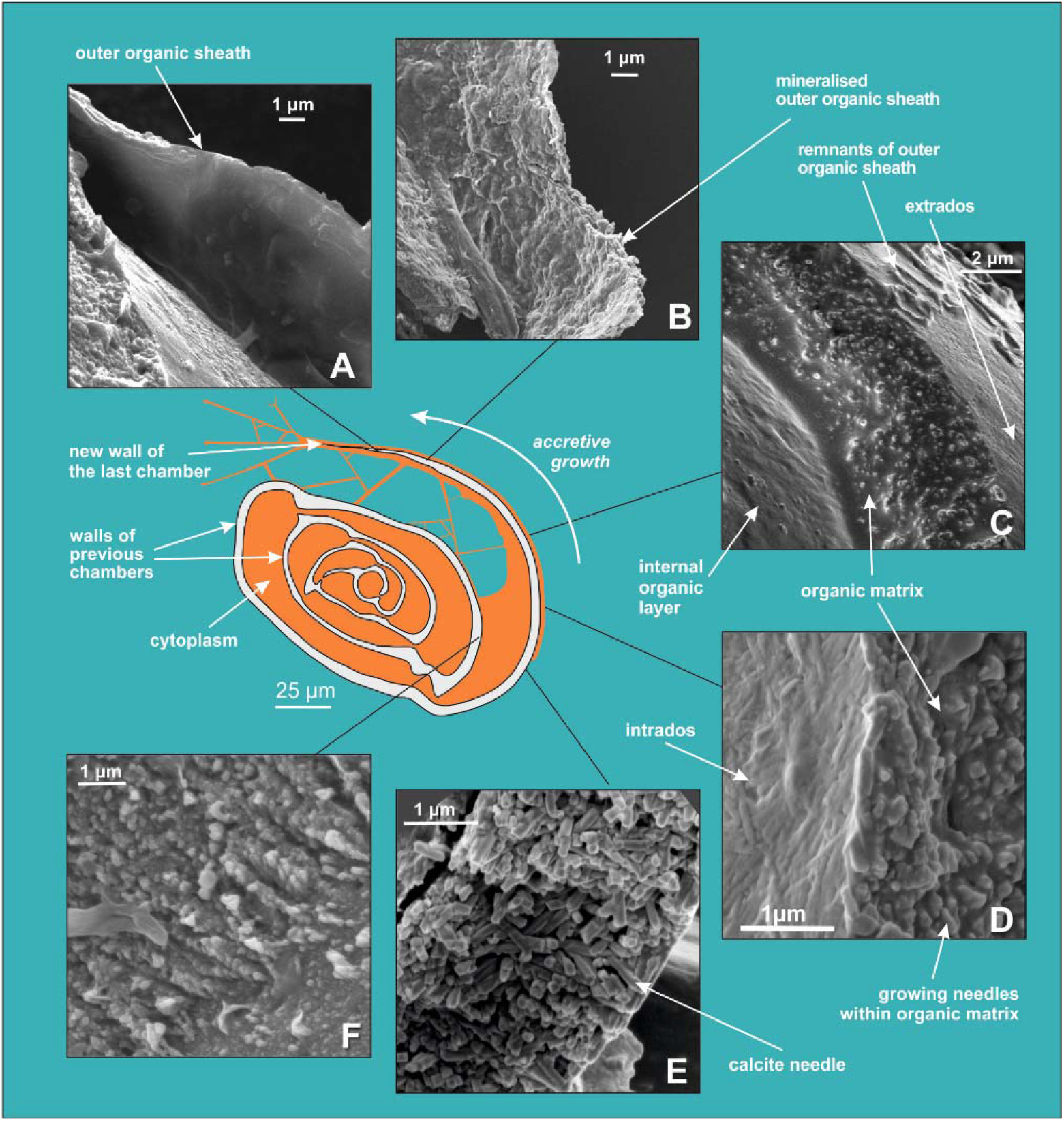
SEM images showing successive stages of new chamber formation in *P. eburnea*. (A) outer organic sheath, (B) mineralized outer organic sheath, (C) calcite nanograins within a gel-like organic matrix, (D) needle-shaped mesocrystal growth, (E) needle-like calcite building elements, (F) nanogranular intrados.

## Discussion

### PORCELANEOUS SHELL FORMATION

Comparative analysis of the nanostructures of the newly built chambers combined with the elemental composition obtained from SEM-EDS, as well as the data from CLSM, allowed us to identify important steps in the accretive formation of *P. eburnea* shells. The formation of a new chamber begins with the construction of a thin outer organic sheath (OOS) that pre-shapes the new chamber (Figs. 4A, 5). The outer organic sheath is made by pseudopodial structures supported by the cytoskeleton immediately after the extrusion of a small mass of cytoplasm from the aperture (Figs. 1E, and 1F). Once the OOS is constructed, the first calcium carbonate accumulation takes place inside in the form of carbonate nanograins (Figs. 4B, 5, S2A and S2B), creating the extrados. The extrados stabilizes the final chamber morphology relatively quickly. Subsequently, the wall gradually thickens through the primary accumulation of hydrated and amorphous Mg-rich CaCO_3_ (Figs. 4B, 5). We suppose that the carbonate content is successively deposited by exocytosis of Mg-ACC rich vesicles that most likely represent the vesicles converted from seawater stained with Calcein (Fig. 5). The characteristic autofluorescence inside foraminiferal cell excited at 405 nm (Fig. 2; Movies S3 and S4) most likely indicates the carbonate content of the vesicles, which are considered to be Mg-ACCs (see Dubicka et al., 2023). Mg-ACC is an unstable, amorphous and hydrated form of CaCO_3_ with a significantly high concentration of Mg (Raz et al., 2000; Weiner et al., 2003; Bentov and Erez, 2006; Kahil et al., 2021) and is commonly regarded as a resource for most biocalcification processes. ACCs have been found in many calcifying marine organisms, such as echinoderms, mollusks, coccolithophorid algae, cyanobacteria, crustaceans, and rotaliid foraminifera, where they are typically interpreted as pre-material phases for the production of calcite skeletons (Hasse, et al., 2000; Weiss et al., 2002; Sviben et al., 2016; Dubicka et al., 2018; Kahil et al., 2021). Research suggests that a high Mg content not only makes ACC unstable but also facilitates the transport of ACC to the crystallization site, where it is initially transformed into carbonate nanograins (Cölfen and Qi, 2001; Addadi and Weiner, 2003; Raz et al., 2003; Dubicka et al., 2023). The existence of intracellular, vesicular intermediate amorphous phase (Mg-ACC pools), which supplies successive doses of carbonate material to shell production, might be supported not only by autofluorescence (excitation at 405 nm; Fig. 2; Movies S3 and S4; see Dubicka et al., 2023) but also by a high content of Ca and Mg analyzed in the cytoplasmic area by SEM-EDS analysis (Fig. S4). In the future, more precise higher resolution elemental measurements are needed for better documentation of miliolid ACC-bearing vesicles. However, the small size of carbonate-bearing vesicles (approximately 1–2 μm) may make this difficult.

**Figure 5.**
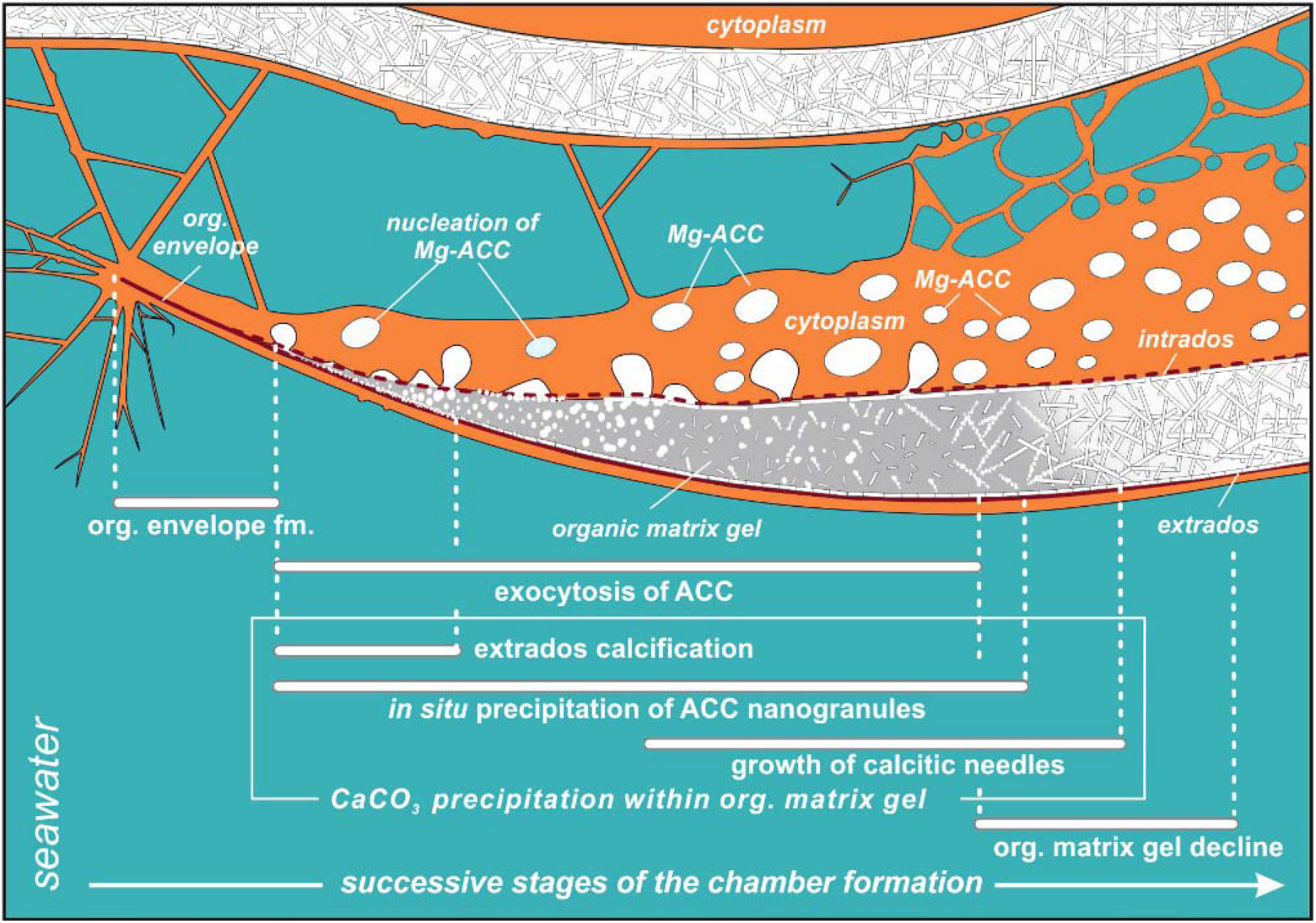
Simplified model of porcelaneous wall construction based on foraminifer *P. eburnea*. White spots labeled as Mg-ACC represent vesicles with Mg-rich amorphous calcium carbonates.

Mg^2+^ and Ca^2+^ ions for intravesicular production of Mg-ACCs are obtained from seawater and taken up by endocytosis, as independently indicated by membrane impermeable Calcein, as well as by the FM1-43 probe selectively labelling membranes of endocytic vesicles (Fig. 2A, B, Movies S1 and S2). We hypothesize that vesicles are carried along cytoskeletal structures to the OM, as observed in rotaliid foraminifera (Dubicka et al., 2023), where they dock and release their contents (Fig. 4). The nanograins then precipitate within the gelatinous matter that consists of amorphous carbonates and organic matrix released from the vesicles (Figs. 3A-C, 4C, and 5). Nanograins immersed in the gel-like matter gradually grow into needle-shaped elements, precipitating *in situ* within the final wall structure (Figs. 3C, 3D, and 4D). The gel-like matter appears to be involved in needle formation; however, the OM seems to disappear (Figs. 3E and 4E) when the needle-shaped crystals are created. We suspect that the gel-like matter consists of pre-formed liquid amorphous mineral phase (Mg-ACC) within the extracellular organic matrix that is suggested by the EDS spectra of the early stage of the wall calcification (Fig. S4: A3 area). The calcification of extrados and intrados occurs before the interior of the wall crystallizes, providing stability to the new chamber at both edges of the wall (Fig. 4D).

The protruding cytoplasm appears to immediately form a chamber wall by secreting OM and crystals from the vesicles (Angell, 1980). As calcite secretion continues along the leading edge, the newly formed segment remains covered by a thin, moving sheet of cytoplasm that is called by Angell (1980) the “active sheet”. This thin active sheet of cytoplasm may represent a lamellipodium that is a pseudopodial structure known to be involved in the biomineralization of Rotaliida (Tyszka et al., 2019). It is also likely that reticulopodial structures (that do not coat the whole calcification site) are responsible for the distribution and shape of the internal surface of the chamber wall. That occurs by successive accumulation of ACC and OM as identified on TEM images by Angell (1980; p. 97). His results suggest that crystallization of calcite needles is “limited to a confined space controlled by active cytoplasmic structures” that are strictly separated by the membranes from the cytosol.

### FORMATION OF SHELL CRYSTALLITES: A PARADIGM SHIFT

Miliolids were thought to share a similar, intracellular, crystallization pathway as the coccolith formation in coccolithophorids (Weiner and Addadi, 2011) that evolved in the Triassic, that is c. 210 Myr ago (Gardin et al., 2012). Coccoliths are produced within intracellular Golgi-derived vesicles and then exported to the surface of the extracellular coccosphere (de Vrind-de Jong et al., 1986). Miliolids, with their unique fibrillar calcitic microstructures, evolved much earlier, that is ca. 300 Myr ago in the late Paleozoic (Fig. S5). Until now, it was generally considered that miliolid crystals also precipitate within vesicles immersed in the cytoplasm and are then transported to the location of the wall construction, where they are released by exocytosis (Berthold, 1976; Angell, 1980; de Noojier et al., 2009; Hemleben et al., 1986; Weiner and Addadi, 2011). Our FE-SEM study of *P. eburnea* shows the lack of premade needle-like crystallites of calcite at the early stages (I-IV) of the wall formation. In contrast, we can clearly infer the *in situ* calcification front with a progressive sequence of crystal growth behind the leading edge of the forming chamber (Figs 4, 5). Therefore, this miliolid species apparently does not produce shells by “agglutination” of premade needle-like crystallites of calcite, in contrast to the traditional miliolid calcification model (Berthold, 1976; Angell, 1980; Hemleben et al., 1986).

In the light of these results, another argument emerges that further confirms in situ calcification of miliolid chambers. It explains the extended transparency of unmineralized walls observed under the light stereomicroscope. The chamber wall under formation tends to gradually change its appearance during calcification from completely transparent to milky and opaque (Fig. 1E, F).

Our results on biomineralization of this miliolid species do not confirm the formation of individual skeletal crystallites within intracellular vesicles. However, in turn, our results do support existence of endocytotic vacuolization of sea water in miliolids that was first suggested by Hemleben et al. (1986). We further support Angell’s (1980) interpretation that the calcite crystals are dispersed in the gel-like organic matrix (see Figs 3A, B; 4C, D; 5). This gel-like fluidal organic matrix most likely include a rich Mg-ACC component as the substrate for *in situ* calcification (Figs 3-5). Interestingly, the previous studies by Angell (1980) did not support crystal formation within vacuoles either.

Precipitation of calcite nanograins, which then merge and transform into crystallites, occurs within the organic matter after the release of Mg^2+^ from Mg-ACC. The organic matter provides an appropriate physiochemical microenvironment for initiating and maintaining the crystallization process by manipulating many essential factors including pH, and kinetics of the system (Kahil et al., 2021). According to Tyszka et al. (2021), the organic matrix involved in the biomineralization of foraminiferal shells may contain collagen-like networks.

Our *in vivo* CLSM observations show a miliolid cytoplasm containing intracellular carbonate-bearing vesicles. Such vesicles have been well-documented by Angell (1980), who stressed their crucial role in the biomineralization process. However, rather than transporting pre-formed solid needles, the vesicles likely carry liquid or quasi-liquid calcification substrates. The liquid phase of the ACC apparently was maintained by a relatively high concentration of Mg (S4), which was much higher than that in the shell, as detected by the SEM-EDS analyses.

Recently, an independent study was performed on another miliolid species - *Sorites orbiculus* (Nagai et al., 2023). The researchers reported highly complementary results that indicate the lack of crystal-like structures within the intracellular vesicles. Their results suggested that calcification of this miliolid species did not follow Hemleben’s et al (1986) model because intracellular vesicles did not produce needle-like crystals to construct the shell wall. They also stated that their observations “may reveal a novel and unknown mode of biomineralisation in foraminifera”.

Because, miliolid wall texture originated together with the appearance of miliolid foraminifera as it has also been recorded within Paleozoic taxa (Fig. S5) thus the calcification mode of miliolids apparently evolved in the late Paleozoic (≥ 350 Mya) and is well conserved in this clade till today. It should be emphasized that our recent understanding of all calcification pathways in Foraminifera implies their independent evolution within main phylogenetic groups, besides miliolids and rotaliids, also including spirillinids, nodosariids, and robertinids (Mikhalevich, 2004; Pawlowski et al., 2013; Mikhalevich, 2014; Dubicka et al., 2018; Dubicka, 2019; Mikhalevich, 2021; Sierra et al., 2022; de Nooijer et al., 2023). In fact, most of these biomineralization evolutionary transitions from agglutination to calcification originated in the mid and late Paleozoic. Mg-ACC has also recently been documented in rotaliid foraminifera (Dubicka et al., 2023). Therefore, the biocalcification processes in Rotaliida and Miliolida, which belong to the two main foraminiferal classes Globothalamea and Tubothalamea, respectively (Pawlowski et al., 2013), are more similar than previously thought (Weiner and Addadi, 2011). Their mesocrystalline chamber walls are created by accumulating and assembling particles of pre-formed liquid amorphous mineral phase. Their calcification occurs within the extracellular organic matrix enclosed in a biologically controlled privileged space by active pseudopodial structures. However, we are aware that this process must also vary to some extent as the chemical composition of the calcite, as well as primary crystallite geometries differ between the groups. Seawater provides the relevant Ca and Mg ions for calcification, which are taken up in both groups by endocytosis. In *Amphistegina* (Rotaliida), this process is performed by shell pores (Dubicka et al., 2023), as well as apertures and foramina; in non-porous Miliolida, it is done by granuloreticulopodia emanating from the aperture (Fig. 2, Movie S2). In both the rotaliid *Amphistegina* and the miliolid *P. eburnea* carbonate-bearing vesicles are surrounded by moving acidic vesicles (Fig. 2, Movies S3 and S4), which likely facilitate pH regulation at the mineralization front (see Toyofuku et al., 2017; Chang et al., 2023). It is very likely that pH is controlled by active outward proton pumping by a V-type H+ ATPase or proton outflux driven by pH that is responsible for the proton flux and related calcification (Toyofuku et al., 2017; see also Matt et al., 2022). We suspect much higher pH values within vesicles transporting Mg-ACC to the site of calcification. Such alkaline vesicles were detected by the HPTS fluorescent labelling and reported by several previous studies (de Nooijer et al., 2008; 2009).

Our findings are in line with recent work in biomineralization, supporting that “biominerals grow by the accretion of amorphous particles, which are later transformed into the corresponding mineral phase” (Macías-Sánchez et al., 2011, p. 1; see also Meldrum and Cölfen, 2008). Miliolid needles, assembled with calcite nanoparticles, are unique examples of biogenic mesocrystals (see Cölfen and Antonietti, 2005), as they form distinct geometric shapes limited by planar crystalline faces. Mesocrystals are constructed from highly ordered individual nanoparticles (Cölfen and Mann, 2003; Strum and Cölfen, 2016; 2017) that form hierarchically structured solid materials in the crystallographic register and are rather devoid of outer planar surfaces. These result from the aggregation, self-assembly, and mesoscopic transformation of amorphous precursor nanoparticles. Mesocrystals are common biogenic components in the skeletons of marine organisms, such as corals, echinoderms, bivalves, sea urchins, and rotaliid foraminifera (e.g., Macías-Sánchez et al., 2011; Benzerara et al., 2011; Seto et al., 2012; Evans et al., 2019; Dubicka et al., 2023).

Our biomineralization model further explains the random orientation pattern of the calcite needles within the shell wall. The miliolid intertwined calcitic structure cannot be explained by the models proposed by Berthold (1976) and followed by Hemleben (1986), that is, by the successive deposition of vesicles with ready bundles of solid calcitic fibers (needles) without additional recrystallization processes. In our proposed *in situ* calcification model, calcite crystallites have sufficient space to grow within the flexible gelatinous organic matrix. In addition, our model explains the need for a light and dark phase for the algae that are present inside *P. eburnea* during the biomineralization processes, as these algae possibly play an important role. Small miliolid coiling foraminifera has been regarded as a non-symbiotic taxon because their shells are not transparent, however, this is not true for red and infrared light. Fully developed miliolid shells are made of randomly distributed needles that cause light reflection, resulting in opaque (porcelaneous) walls that possibly protect the foraminifera from UV irradiation and allow them to live in extremely illuminated shallow seas (Hohenegger, 2009). These walls are permeable to red and infrared light, as we observed using multiphoton laser. Red light is commonly believed to be the most efficient waveband for photosynthesis however green light may achieve higher quantum yield of CO_2_ assimilation and net CO_2_ assimilation rate (Liu and Lerser, 2021). *Pseudolachlanella eburnea* may acquire its facultative symbionts only for the duration of the biomineralization process. The late stage of needle formation in the shell production process ensures that the wall remains transparent by the time the needles are completed. Similar patterns of the gradual change from transparent to opaque whitish walls were also observed in larger symbiotic miliolids by Marshalek (1969), Wetmore (1999), and Tremblin et al. (2022). The latter authors (Tremblin et al., 2022) documented chamber formation of miliolid *Vertebralina striata* with cytoplasm enveloped by a transparent sheath decorated with striate already present in the transparent wall before calcification. They also interpret white areas on the sheath, indicating incipient concentrations of minute calcite crystallites that represent the mineralised wall. The biomineralization process is likely aided by their dark respiratory activity (see Hallock, 1999), as they could supply calcification substrates such as HCO^3-^ through respiration or by increasing pH at the calcification site during the light phase. Similarly, representatives of miliolid large benthic foraminifera (Archaiasidae, Soritidae, and Peneroplidae) host endosymbiotic algae (Lee, 2006; Prazes and Renema, 2019). Therefore, they have developed additional morphological and textural features such as pits, grooves/striate, or windows, which enable light penetration into the places where symbionts are positioned (see Hohenegger, 2009, Parker 2017).

## Materials and Methods

Living foraminifera, collected from the coral reef aquarium in the Burgers’ Zoo (Arnhem, Netherlands), were cultured in a 10 L aquarium containing seawater with a salinity of 32‰, pH of 8.2, and a temperature of 24 °C. *Pseudolachlanella eburnea* (d’Orbigny) was placed in 4 mL petri dishes one day before CLSM studies and observed under a Zeiss Stemi SV8 Stereomikroskope. Selected individuals were studied *in vivo* using a Leica SP5 Confocal Laser Scanning Microscope equipped with an argon, helium-neon, neon, diode, and multiphoton Mai Tai laser (Spectra-Physics) at the Alfred-Wegener-Institut, Bremerhaven, Germany. In *vivo* experiments were performed by labeling samples with different fluorescent dyes (Table 1) just before imaging using, p*H-*sensitive LysoGlow84 (50 μM exc. MP720 nm exc/em. 380–415 nm and 450–470 nm, Marnas Biochemicals Bremerhaven, incubation time: 2 h), FM1-43 membrane stain (1 μM, exc. 488 nm em. 580–620 nm, Invitrogen, incubation time: 24 h), and membrane impermeable Calcein (0.7 mg/10 mL, exc. 488 nm, em. 510–555 nm, incubation time: 24 h). The foraminifera were removed from the Petri dish with clean water using a pipette. In addition, the autofluorescence of specific foraminiferal structures at the chosen excitation/emission wavelength was detected. All experiments were replicated with at least several individuals of the same species. All fluorescence probe experiments were performed with appropriate controls.

**Table 1.**
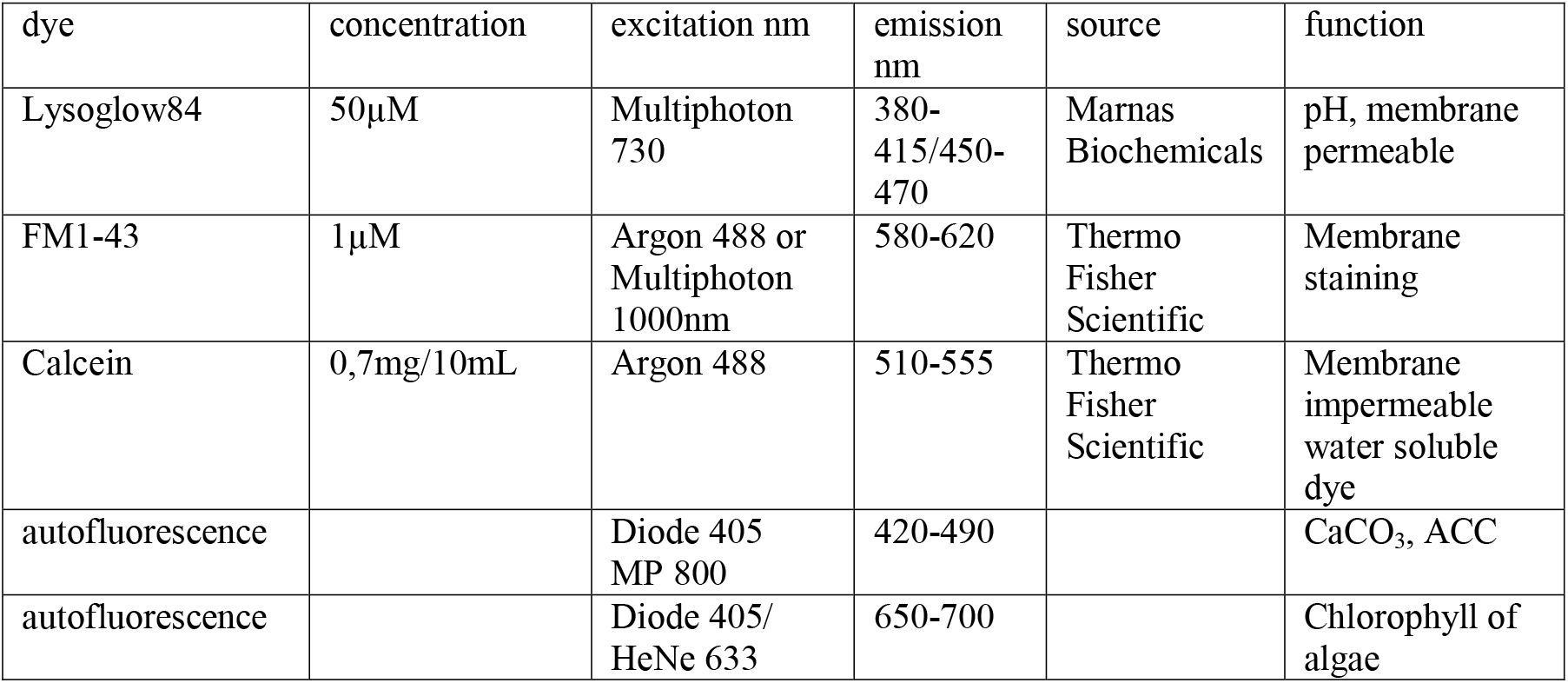
Wavelengths and dyes.

| dye | concentration | excitation nm | emission nm | source | function |
| --- | --- | --- | --- | --- | --- |
| Lysoglow84 | 50µM | Multiphoton 730 | 380-415/450-470 | Marnas Biochemicals | pH, membrane permeable |
| FM1-43 | 1µM | Argon 488 or Multiphoton 1000nm | 580-620 | Thermo Fisher Scientific | Membrane staining |
| Calcein | 0,7mg/10mL | Argon 488 | 510-555 | Thermo Fisher Scientific | Membrane impermeable water soluble dye |
| autofluorescence |  | Diode 405 MP 800 | 420-490 |  | CaCO<sub>3</sub>, ACC |
| autofluorescence |  | Diode 405/ HeNe 633 | 650-700 |  | Chlorophyll of algae |

Additional foraminifera individuals that had been studied by CLSM were fixed for further analysis. The fixation process followed two different methods: (A) 60 individuals were transferred to 3% glutaraldehyde for 5 s and then dehydrated stepwise for a few seconds with an ethanol/distilled water mixture with increasing concentrations (30%, 50%, 70%, and 99%). (B) The seawater was removed from 50 individuals by pipetting and applying a small piece of Kimtech lab wipe (without any rinsing), followed by quick drying in warm air (30–35 °C). This method stops the dissolution of the amorphous mineral phase because there is no contact with other liquids. Fixed foraminifers of both procedures were gently broken using a fine needle to coat the cross-sectional surfaces and tested inside with a few nanometers of either gold or carbon. Foraminifera were then studied using a Zeiss Σigma variable-pressure field-emission scanning electron microscope (VP-FESEM) equipped with EDS at the Faculty of Geology, University of Warsaw.

## Acknowledgments

This work was supported by the Alexander von Humboldt Foundation Research Fellowship for experienced researchers to Z.D. and the Polish National Science Center (UMO-2018/29/B/ST10/01811) to J.T. and Z.D., a grant coordinated by Grzegorz Racki, University of Silesia. We thank Oscar Branson for his comments on an earlier version of the manuscript and the suggested improvements.

## Movie captions

**Movie S1 (separate file)**. Living *P. eburnea* showing cell impermeable Calcein (blue, exc. 488nm em. 505-555) in a series of 107 overlaid images taken during 428 s. Calcein staining indicates the occurrence of seawater vesicles inside the cytosol.

**Movie S2 (separate file)**. FM1-43 membrane probe fluorescent signals (red, exc. 488nm, em. 580-620nm) emitted by intracellular vesicles within cytosol of *P. eburnea*. Because FM1-43 stains only the cell membrane, the observed vesicles must be originated during the process of endocytosis. The movie was taken by overlaid of 84 images during 433 s.

**Movie S3 (separate file)**. Living *P. eburnea* showing fluorescence signal inside the cytosol: autofluorescence of Mg-ACC pools (red, exc. 405nm, em. 420-490nm) and algal chloroplasts (green, exc. 633nm, em. 640-690nm), fluorescent signal of LysoGlow84 pH sensitive dye (exc. MP 720nm, em. 440-470nm) indicating acidic vesicles. The movie was taken by overlaid of 37 images during 555 s.

**Movie S4 (separate file)**. Living *P. eburnea* showing fluorescence signal inside the cytosol: autofluorescence of Mg-ACC pools (red, exc. 405nm, em. 420-490nm) and algal chloroplasts (green, exc. 633nm, em. 640-690nm), fluorescent signal of LysoGlow84 pH sensitive dye (exc. MP 720nm, em. 440-470nm) indicating acidic vesicles. The movie was taken by overlaid of 37 images during 555 s.

## Notes

### Competing Interest Statement

The authors have declared no competing interest.

### Summary of Updates

We toned down the the physiological interpretation based on fluorescence studies in the revised version of the manuscript.

