## Supplemental figures for "Biocalcification in porcelaneous foraminifera"

SUPPLEMENTARY FIGURES

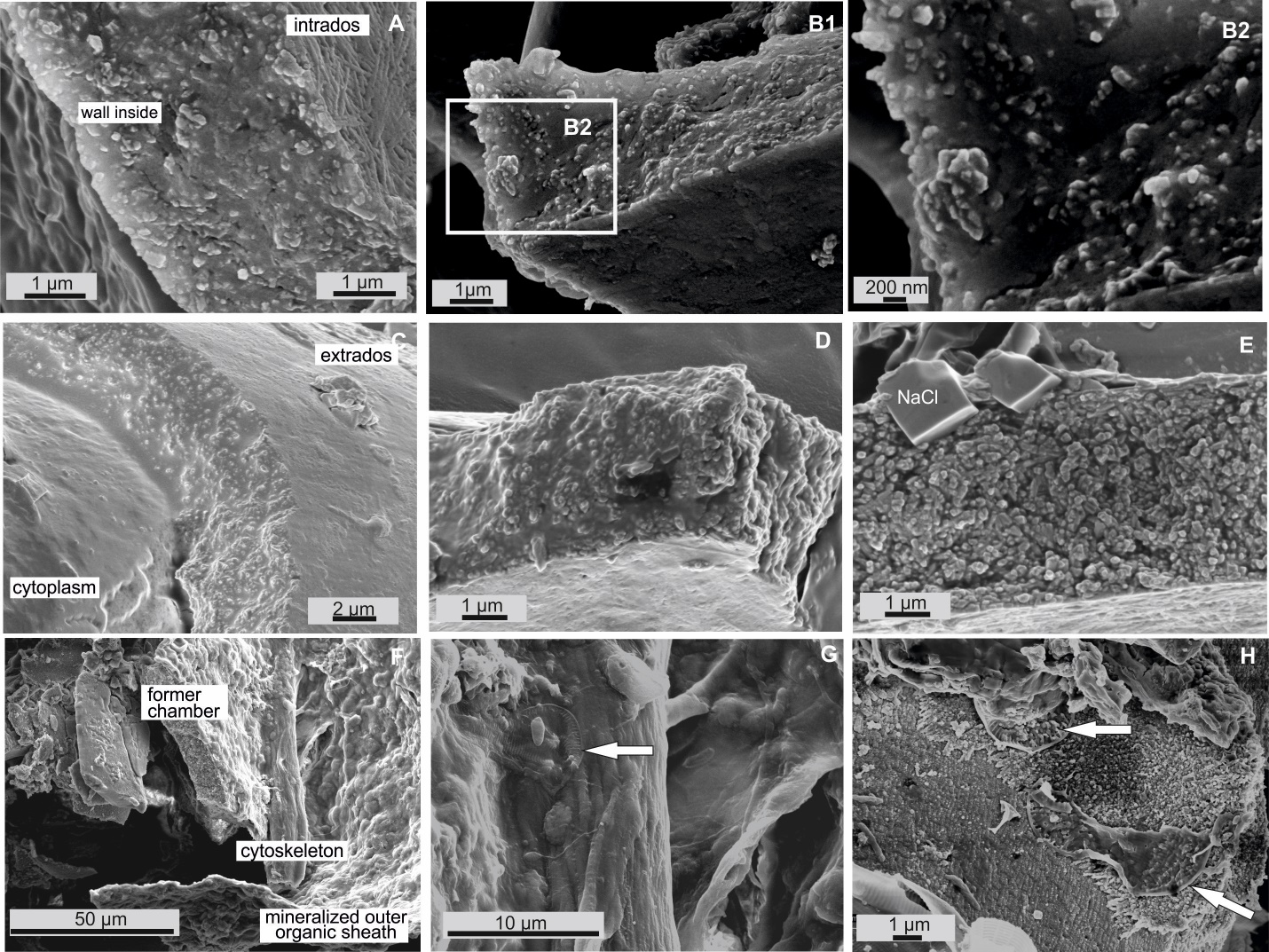

**Figure S1.** SEM images of broken specimen of *Pseudolachlanella eburnea* (d’Orbigny) showing the test wall made of calcite nanograins (A-E) within organic gel-like fluids in-between, and cytoplasmic structures below a newly created chamber (G), and diatoms (white arrows) inside the test (G, H).

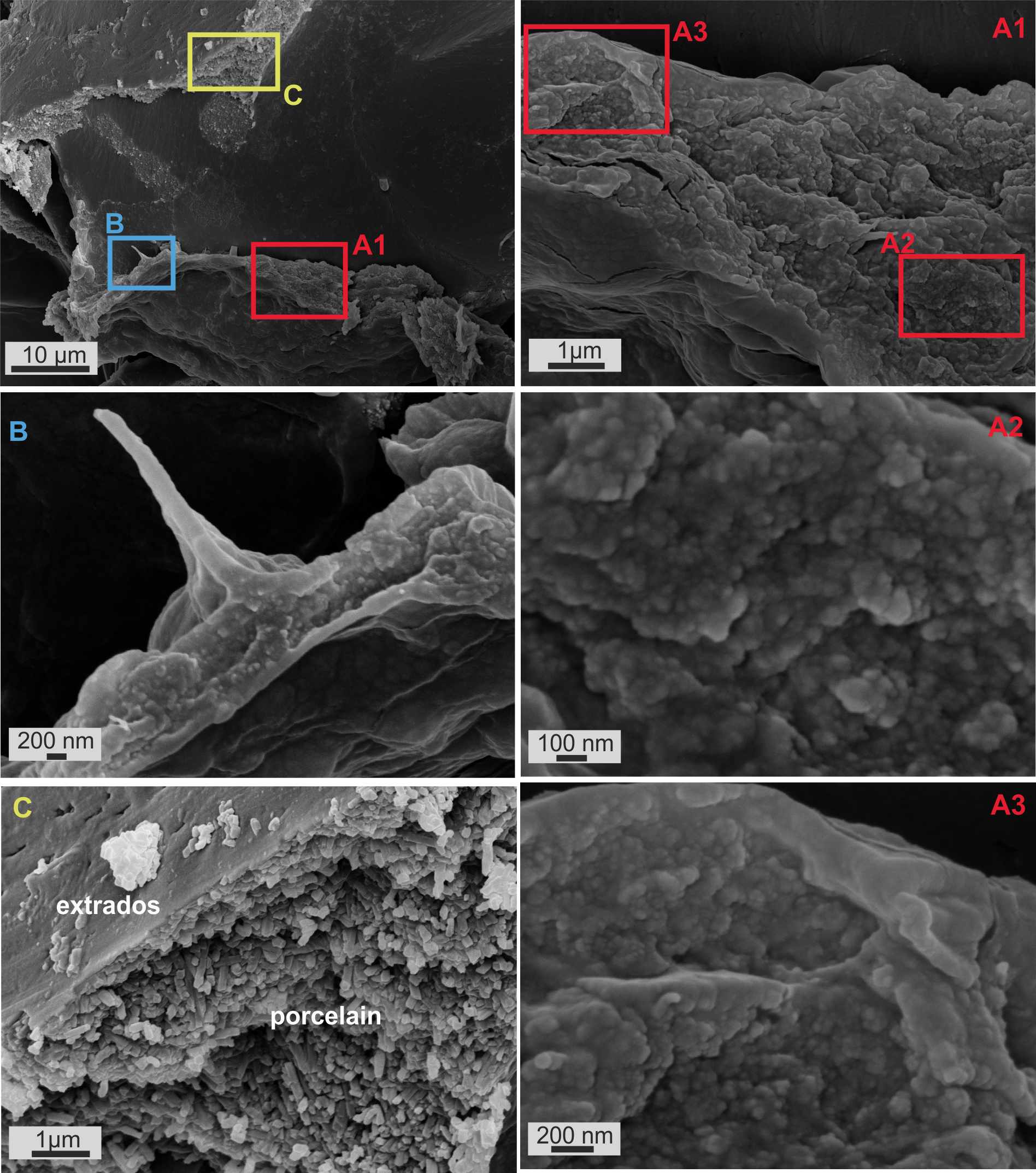

Figure S2. SEM images of broken specimen of *Pseudolachlanella eburnea* (d’Orbigny) showing a cross-section of the mineralized outer organic sheath of the last chamber (A, B) and the fully mineralized test wall of the former chamber (C).

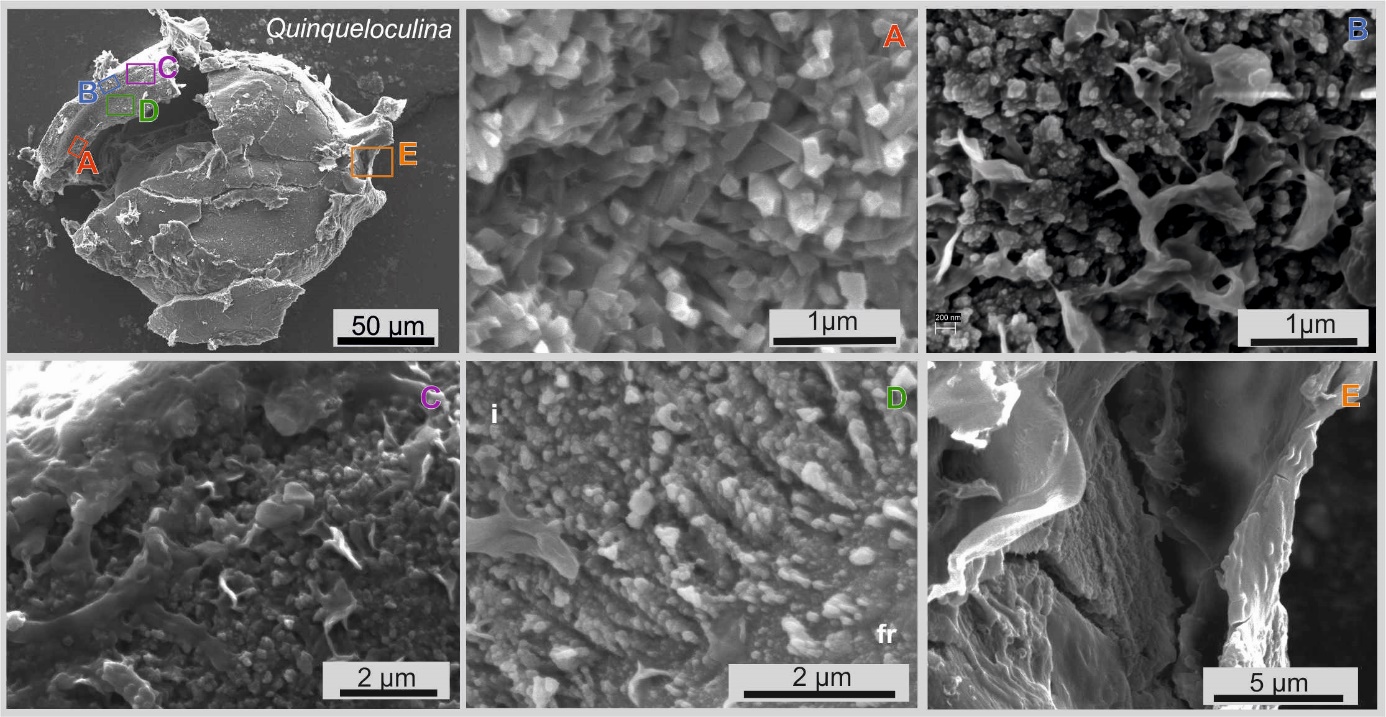

Figure S3. SEM images of different parts of the broken test of the *Pseudolachlanella eburnea* (d’Orbigny) specimen showing (A) a fully developed test wall made of randomly oriented calcite needle-shaped crystals with planar faces; (B) test inside of newly built chamber wall made of calcite nanograins and some organic matter between the needles; (C) test inside of newly built chamber wall covered with an outer organic sheath; (D) rudiments of needles of extrados composed of rows of calcite nanograins and attached to the former chamber (fr); (E) outer organic sheath.

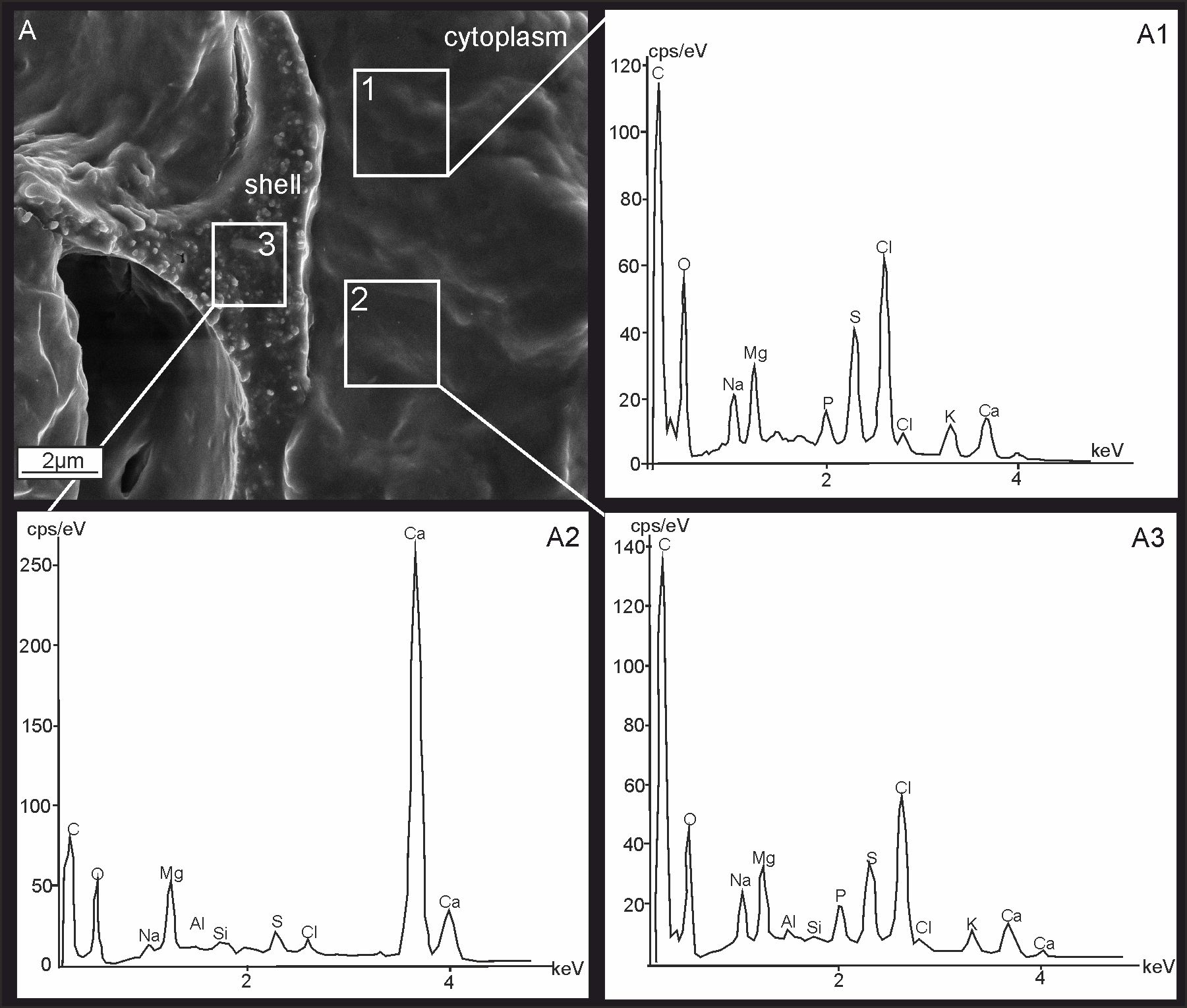

**Figure S4.** SEM images (A) and EDS spectra (A1-A3) of *P. eburnea* fixed cytosol (A1, A2) and newly created chamber (A3) indicating the chemical composition of both structures. cps/eV: counts per second per electron-volt.

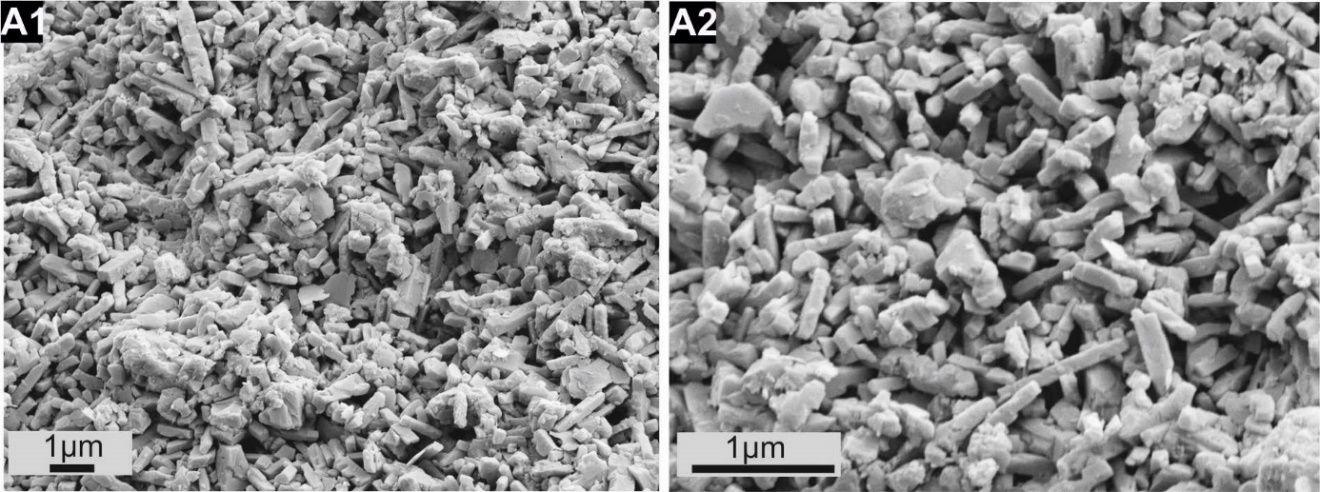

**Figure S5.** SEM images of miliolid *Agathamina pusilla* Geinitz from the lower Permian (ca. 290 Mya) of the Holy Cross Mountains (Poland) showing needle test structure identical to that of Recent taxa.

Movie S1 (separate file). Living *P. eburnea* showing cell impermeable Calcein (blue, exc. 488nm em. 505-555 nm) in a series of 107 overlaid images taken during 428 s. Calcein staining indicates the occurrence of seawater vesicles inside the cytosol.

Movie S2 (separate file). FM1-43 membrane probe fluorescent signals (red, exc. 488nm, em. 580-620nm) emitted by intracellular vesicles within cytosol of *P. eburnea.* Because FM1-43 stains only the cell membrane*,* the observed vesicles must be originated during the process of endocytosis. The movie was taken by overlaid of 84 images during 433 s.

Movie S3 (separate file). Living *P. eburnea* showing fluorescence signal inside the cytosol: autofluorescence of Mg-ACC pools (red, exc. 405nm, em. 420-490nm) and symbiontic chloroplasts (green, exc. 633nm, em. 640-690nm), fluorescent signal of LysoGlow84 pH sensitive dye (exc. MP 720nm, em. 440-470nm) indicating acidic vesicles. The movie was taken by overlaid of 37 images during 555 s.

Movie S4 (separate file). Living *P. eburnea* showing fluorescence signal inside the cytosol: autofluorescence of Mg-ACC pools (red, exc. 405nm, em. 420-490nm) and chloroplasts (green, exc. 633nm, em. 640-690nm) of microalgae, fluorescent signal of LysoGlow84 pH sensitive dye (exc. MP 720nm, em. 440-470nm) indicating acidic vesicles. The movie was taken by overlaid of 37 images during 555 s.
